# SDC4 deletion perturbs intervertebral disc matrix homeostasis and promotes early osteopenia in the aging spine

**DOI:** 10.1101/2023.09.24.559195

**Authors:** Kimheak Sao, Makarand V. Risbud

## Abstract

Syndecan 4 (SDC4), a cell surface heparan sulfate proteoglycan, is known to regulate matrix catabolism by nucleus pulposus cells in an inflammatory milieu. However, the role of SDC4 in the aging spine has never been explored. Here we analyzed the spinal phenotype of SDC4 global knockout (KO) mice as a function of age. Micro-computed tomography showed that SDC4 deletion severely reduced vertebral trabecular and cortical bone mass, and biomechanical properties of vertebrae were significantly altered in SDC4 KO mice. These changes in vertebral bone were due to elevated osteoclastic activity in KO mice. The histological assessment also showed subtle phenotypic changes in the intervertebral discs. Imaging-Fourier transform-infrared (FTIR) analyses showed a reduced relative ratio of mature collagen crosslink in young adult NP and AF compartments of KO compared to wildtype (WT) mice. Additionally, relative chondroitin sulfated glycosaminoglycan (GAG) levels increased in the NP compartment of the KO mice. Transcriptomic analysis of NP tissue using CompBio, an AI-based tool showed biological themes associated with prominent dysregulation of heparan sulfate GAG degradation, mitochondria metabolism, autophagy, and endoplasmic reticulum to Golgi protein processing. Overall, this study highlights the important role of SDC4 in fine-tuning vertebral bone homeostasis and extracellular matrix homeostasis in the intervertebral disc.

## Introduction

Low back pain is highly prevalent and the number one leading cause of years lived with disability worldwide(1). Although the etiology of low back pain is multifactorial, age-associated intervertebral disc degeneration is one major contributor to this pathology(2). The disc confers flexibility, permits motion, and transmits applied mechanical loads to the spinal column(3). This is due to three unique compartments: the avascular, gelatinous core nucleus pulposus (NP) concentrically constrained by a fibrocartilaginous annulus fibrosus (AF) and both capped superiorly and inferiorly by cartilaginous endplates covering the contiguous vertebra. The hypoxic NP compartment is rich in aggrecan proteoglycans. Their negatively charged glycosaminoglycan (GAG) side chains draw water within the extracellular matrix (ECM) to hydrate the disc(4). In aging, gradual breakdown and loss of PG molecules compromise mechanical function of intervertebral discs promoting degeneration(5, 6). Of particular interest is heparan sulfate proteoglycan (HSPG) syndecan 4 (SDC4), known for its diverse role in modulating cell signaling via binding of soluble factors, cell-associated molecules, or ECM components(7, 8).

SDC4 is a cell surface proteoglycan that mediates numerous cellular processes that affect proliferation, differentiation, mechanotransduction via focal adhesion formation and matrix catabolism(9–13). SDC4 knockout (KO) mice develop impaired responses to tissue injury and bone fracture repair(14, 15). In osteoarthritis (OA), SDC4 regulates breakdown of cartilage via aggrecanase, disintegrin and metalloproteinase with thrombospondin type I motif, member 5 (ADAMTS-5) through mitogen-activated protein kinase (MAPK)-dependent-matrix-metalloproteinase 3 (MMP3) signaling axis(16). In the NP tissue, we have shown that SDC4 promotes cytokine-dependent activity of ADAMTS-5 independent of MMP3 and contributes to Sox9 modulation(16, 17). However, the role of SDC4 in the aging spine has not been investigated *in vivo*. Herein we have characterized age-dependent changes in the motion segment of SDC4 KO mice. Since a pathological hallmark of disc degeneration is the imbalance between production of catabolic and anabolic factors by resident cells, we specifically examined SDC4’s role in matrix homeostasis. Importantly, low back pain is correlated with the health and integrity of the motion segment comprising both the vertebrae and intervertebral disc(6). The present study provides the first evidence that SDC4 affects the balance between bone formation and resorption in the lumbar vertebrae and ECM maintenance in the disc. Overall, we found striking alterations in vertebra bone geometry and mechanical properties due to the imbalance of osteoblastic-osteoclastic activities and reduction in disc matrix turnover highlighting the importance of SDC4 in the spinal column.

## Results

### SDC4 loss causes early vertebral osteopenia

Previous work has implicated SDC4 in long bone fracture healing(15). To assess the impact of SDC4 global deletion on the vertebral bone during aging, we performed micro-computed tomography (µCT) on the lumbar vertebra (L1-6) of mice at 6-month (young adult), 12-month (middle-age), and 24-month (old) mice. Three-dimensional (3D) reconstruction of vertebra trabecular tissue notably highlighted bone loss during the natural aging in wildtype (WT) mice, but more strikingly, SDC4 deletion severely affected trabecular bone architecture in young adult, middle-aged and old mice (Fig. 1A). µCT analysis revealed that there was significant reduction in bone mineral density (BMD), percent bone volume (BV/TV), trabecular thickness (Tb.Th.) and number (Tb.N.), with a concomitant increase in trabecular separation (Tb.Sp.) in SDC4 KO mice (Fig. 1B-F) at all timepoints. Similar to trabecular tissue, cortical lumbar vertebrae bone was also affected in SDC4 KO mice (Fig. 1G-K). The analysis showed a reduction in tissue mineral density (TMD), cross-sectional thickness (Cs.Th.), mean total cross-sectional bone area (B.Ar.) and tissue area (T.Ar.) in SDC4 KO mice (Fig. 1H-K). The vertebral body length, disc height, and disc height index however did not show significant changes between KO and WT mice (Fig. 1L-N). These changes in vertebral bone health parameters suggested that SDC4 plays an important role in the regulation of trabecular and cortical vertebral bone geometry.

**Fig. 1.**
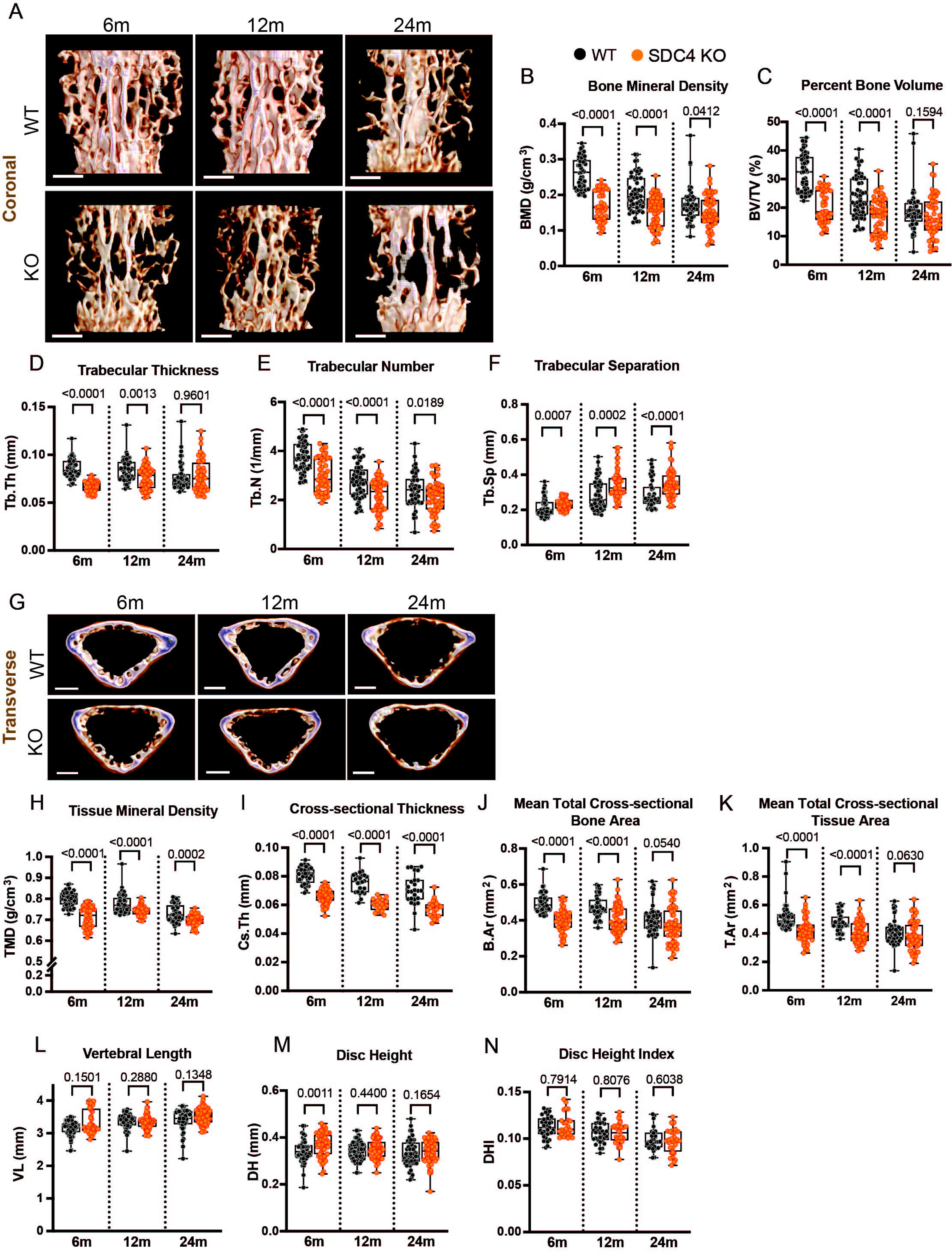
Loss of SDC4 causes reduced vertebral bone mass and altered bone geometry. (A) The representative 3D rendered trabecular lumbar vertebral bone at 6-month, 12-month, and 24-month, showing bone loss in SDC4 KO compared to WT control. Scale bars, 0.5 mm. (B) Bone mineral density (BMD) (g/cm^3^), (C) Percent bone volume/ tissue volume (BV/TV) (%), (D) trabecular thickness (Tb.Th) (mm), (E) trabecular number (Tb.N) (1/mm), (F) trabecular separation (Tb.Sp) (mm) were analyzed. (G) Representative 3D rendered cortical tissue shows thinning of the cortical bone, assessed by (H) Tissue mineral density (TMD) (g/cm^3^), cross-sectional thickness (Cs.Th) (mm), mean total cross-sectional thickness bone area (B.Ar) (mm^2^), mean total cross-sectional tissue area (T.Ar) (mm^2^). (L) Vertebral length (VL) (mm), disc height (DH) (mm), and disc height index (DHI) were also measured. 6-month WT N = 11 mice, n = 55 vertebrae, KO = 11 mice, n = 48 vertebrae; 12-month WT N = 11 animals, n = 54 vertebrae, KO = 11, n = 55 vertebrae; 24-month WT N = 10 animals, n = 50 vertebrae, KO = 10, n = 52 vertebrae. Quantitative measurements represent median with interquartile range. Significance was determined using an unpaired Welch’s t-test or Mann-Whitney test, as appropriate.

### SDC4-loss promotes increased osteoclastic activity in vertebrae

To examine whether altered vertebral bone geometry in SDC4 KO mice was due to an imbalance in activity between osteoblasts and osteoclasts, tissue nonspecific alkaline phosphatase (TNAP) and tartrate-resistant acid phosphatase (TRAP) staining were performed on mineralized tissue sections, respectively. Given that SDC4 KO vertebral bone remodeling is seen at 6-months of age, vertebral sections of young adult mice were used for TNAP and TRAP staining (Fig. 2A). We found that SDC4 KO vertebrae had a moderate reduction in TNAP staining (Fig. 2B) and an increase in overall TRAP staining (Fig. 2C). To further identify if the reduction in TNAP staining seen in SDC4 KO was due to an intrinsic osteoblast differentiation defect, bone marrow-derived stromal cells (BMSCs) from both genotypes were cultured and differentiated into osteoblasts. Alkaline phosphatase and alizarin red staining revealed no obvious defects in osteoblast differentiation and calcium hydroxyapatite crystals deposition (Fig. 2D), respectively in SDC4 KO mice. Interestingly, SDC4 deletion promoted increased osteoclastogenesis as evidenced by increased average osteoclast numbers per well (Fig. 2E-F). These results suggested that the upregulated osteoclastic activity in SDC4 deficient vertebra, and not the dysregulation in osteoblastic activity, was the driver of morphometric deficits we observed in µCT analysis. Altogether, high bone turnover evidenced by elevated osteoclastic activity suggests that loss of SDC4 could promote early osteopenia.

**Fig. 2.**
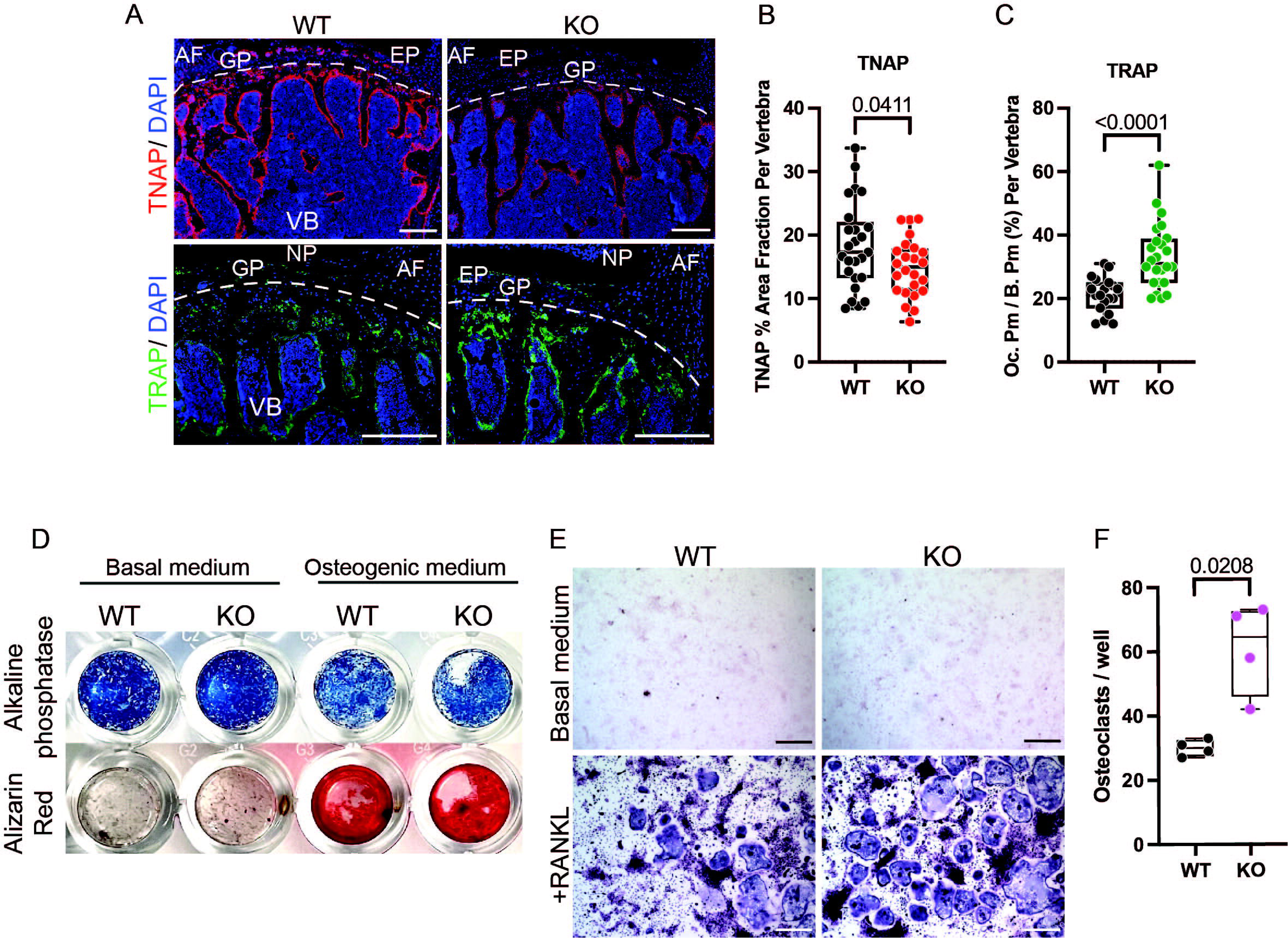
Osteoblastic activity is dysregulated in lumbar vertebrae of SDC4 KO mice. (A-C) Immunohistological staining showed decreased staining of (A, B) TNAP and increased staining of (A, C) TRAP in 6-month SDC4 KO lumbar vertebrae. Scale bars, 500µm, 250µm, respectively. TNAP: WT N = 8 mice, n = 25 vertebrae; KO N = 8, n = 24. TRAP: WT N = 8, n = 19; KO N = 10, n = 24. (D) Osteoblastogenesis of BMSCs shows no difference in mineral deposition between KO and WT mice. N = 2-3 animals/genotype were pooled per experiment. (E, F) Osteoclastogenesis shows an increase in osteoclasts count. Scale bar, 1mm. N = 3-4 animals/genotype were pooled per experiment. Quantitative measurements represent the median with the interquartile range. Significance was determined using an unpaired Welch’s t-test or Mann-Whitney test, as appropriate.

### SDC4 deletion alters vertebral mechanical properties

Given pronounced alterations in vertebral bone geometry in SDC4 KO mice, we next examined the biomechanical properties of KO and WT vertebrae. We performed a monotonic displacement compression test where vertebral bodies were subjected to a 0.4 Newton (N) compressive pre-load, followed by a monotonic displacement ramp at 0.1 mm/s until failure (Fig. 3A). Bone compression results revealed that SDC4-deficient vertebrae had significantly reduced stiffness and Young’s Modulus compared to WT control (Fig. 3B-C), demonstrating that low BMD affects bone stiffness and offers lower resistance to deformation under an applied compressive force. Interestingly, while both WT and KO have comparable intrinsic vertebral bone strength and can withstand similar force to fracture as evidenced by ultimate stress and ultimate load (Fig. 3D-E), KO vertebrae had increased ultimate strain, ultimate displacement, ultimate energy, toughness, and post-yield displacement, suggesting that these vertebrae are more ductile and able to absorb more energy before fracture (Fig. 3F-J).

**Fig. 3.**
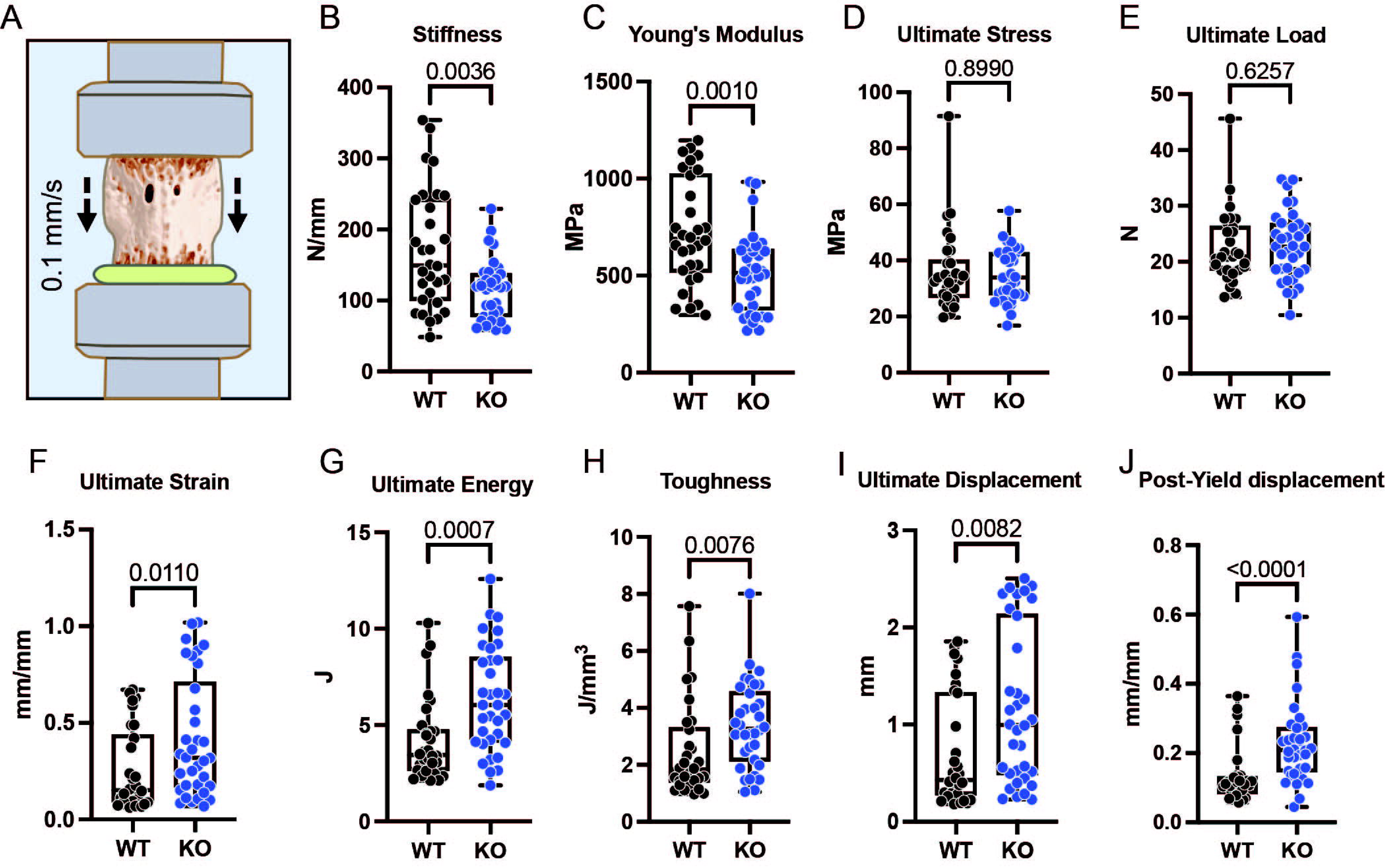
Loss of SDC4 alters mechanical properties of lumbar vertebral bone. (A) Illustration of potted lumbar vertebra under compressive pre-load, followed by a monotonic displacement ramp at 0.1 mm/s until failure. (B) Stiffness (N/mm), (C) Young’s Modulus (MPa), (D) ultimate stress (MPa), (E) ultimate load (N), (F) ultimate strain, (G) ultimate energy (J), (H) toughness (J/mm^3^), (I) ultimate displacement (mm), and (J) post-yield displacement measurements were done to assess the structure and elastic properties of the bones. WT N = 10 mice, n = 30 vertebrae; KO N = 11 mice, n = 34 vertebrae. Quantitative measurements represent the median with the interquartile range. Significance was determined using an unpaired Welch’s t-test or Mann-Whitney test, as appropriate.

### SDC4 deletion causes subtle phenotypic changes in the disc

Previously, we had demonstrated that proinflammatory cytokines TNF-α and IL-1β elevate SDC4 expression in NP cells and that it selectively promotes aggrecan degradation via ADAMTS5(13). Since the positive feedback interplay between SDC4 and proinflammatory cytokines promote catabolic degradation of matrix proteins in the disc, it is of interest to ascertain what role SDC4 plays during disc aging. With respect to the spine aging, Beckett et al. showed that SDC4 expression is highly enriched in the rat intervertebral disc up to 24-months of age, specifically in NP cells(18). We therefore assessed the disc phenotype in SDC4 KO and WT control mice up to 24-months. Histological assessment of 6-month KO and WT lumbar discs using Safranin-O/ Fast Green/ Hematoxylin staining showed that approximately twice the number of SDC4 KO discs scored lower on the average Modified Thompson score of degeneration than their age-matched control discs (Fig. 4A-D). While 6-month young animals are healthy, it is worth noting that SDC4 KO NP compartment showed a significantly thicker NP cell band and increased cell number without altering the NP compartment area or NP aspect ratio compared to WT (Fig. 4A’, E). Interestingly, however, at 12-, and 24-months no appreciable differences in the NP and AF compartment morphology were noted between KO and WT lumbar discs, suggesting that in the context of SDC4 loss, the aging was a predominant determinant of disc health than the genotype. SDC4 is involved in many biological processes, some of which regulates cellular contractility, adhesion, and cytoskeletal organization(19). A recent study by Chronopoulos et al. revealed that SDC4 is a key cellular mechanotransducer that tunes cell mechanics in response to localized mechanical tension and that SDC4-deficient MEFs lacked F-actin stress fibers with low nuclear-to-cytoplasmic YAP ratio due to low cytoskeletal tension(12). To examine if this mechanosensing pathway is dysregulated in SDC4-KO mouse NP (mNP) cells, we cultured freshly isolated SDC4 null and WT mNP cells onto collagen-or fibronectin-coated glass and assessed focal adhesion formation (Fig. 4F-F”). We investigated mechanotransduction ability of these cells using phosphorylated paxillin and YAP staining, respectively. Morphologically, both WT and SDC4 KO mNP cells were able to adhere, spread, and form similar focal adhesion patterns and F-actin stress fibers on both substrates and showed persistence of cytoplasmic vacuoles, outlined by the structurally perturbed F-actin and void of cytoplasmic YAP (Fig. 4F-F”). Interestingly, in contrast to Chronopoulos et al., SDC4-deficient mNP cells were able to sense mechanical tension from the extracellular environment and biochemically relayed that signal to promote nuclear localization of YAP on both substrates (Fig. 4F’’). Together, these results suggest that SDC4 may not be an essential component of a mechanosensory apparatus in mNP cells.

**Fig. 4.**
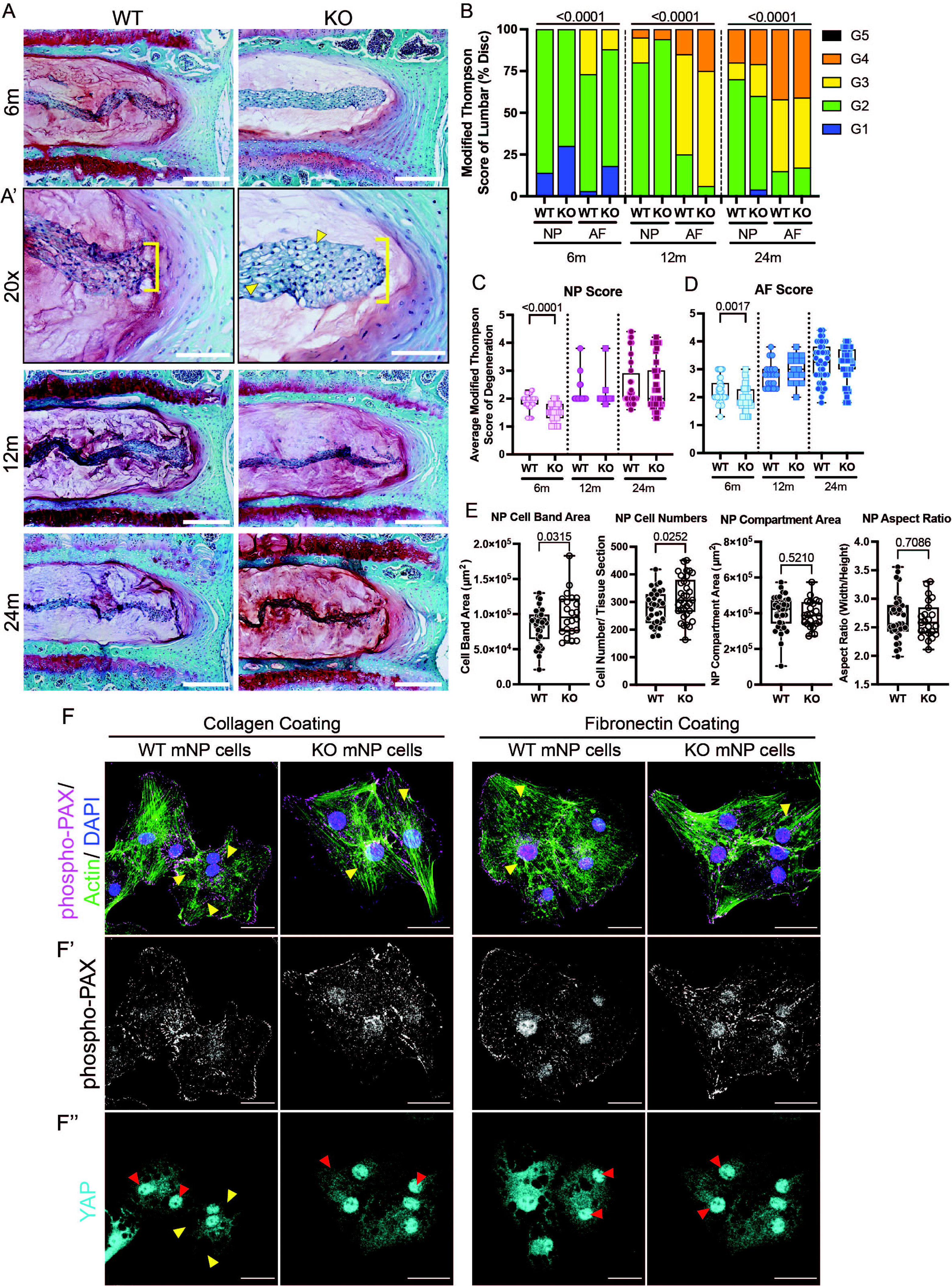
SDC4 deletion causes mild phenotypic changes in the NP compartment of intervertebral disc. (A) Intervertebral discs of WT and KO were assessed using Safranin O/Fast Green staining, showing tissue morphology and proteoglycan content. (B-D) Modified Thompson Grading was used to assess the NP and AF degeneration scores. Scale bars, 250µm. 6-month WT N = 9 mice, n = 37 discs; KO N = 11, n = 40; 12-month WT N = 5, n = 20; KO N = 4, n = 16; 24-month WT N = 10, n = 40; KO N = 12, n = 48. (A’) 6-month discs show increased retention of vacuolated NP cells (yellow arrow heads) and thicker NP cell band (yellow bracket). Scar bars, 100µm. (E) NP cell band area, cell numbers, compartment, and aspect ratio were measured. WT N = 9-11 animals, n = 32-35 discs; KO N = 9-10 mice, n = 23-34 discs. (F) Cell spreading was also assessed on collagen and fibronectin coating using phalloidin and p-PAX. (F’-F’’) Adhesion formation and mechanotransduction of mNP cells were observed using p-PAX and nuclear translocation of YAP (red arrowhead). Scale bars, 25µm. Quantitative measurements represent the median with the interquartile range. Significance was determined using an unpaired Welch’s t-test or Mann-Whitney test, as appropriate.

### SDC4 deletion results in alteration in collagen crosslinks

To examine changes in the collagen I-rich AF compartment of KO discs, we combined picrosirius red staining and polarized microscopy to visualize collagen fiber structure and thickness in young and old lumbar discs (Fig. 5A). Analysis showed increase of red birefringence with concomitant decrease of green birefringence in the KO discs at 6-months (Fig. 5B, D), with no differences noted at 24-months between genotypes (Fig. 5C). We then used Fourier transform infrared (FTIR) spectroscopy imaging to define matrix compositional changes in lumbar discs of young and old mice. The discs were imaged, and second derivative function was used to enhance the separation of overlapping peaks before extracting individual spectra of interest for analysis (Fig. 5E-F), specifically, peaks centered around 1660 cm^-1^ (amide I, pyridinoline [PYR]), 1690 cm^-1^ (dehydro-dihydroxynorleucine [de-DHLNL]), 1064 cm^-1^ (chondroitin sulfated GAG specific peak), 1338 cm^-1^ (collagen specific peak), and 1549 cm^-1^ (amide II)(20) were analyzed. To determine the maturity of collagen fibril, peak height value of non-reducible mature trivalent crosslinks (PYR) at 1660 cm^-1^ was compared to peak height value of the immature divalent PYR precursor crosslinks (de-DHLNL) at 1690 cm^-1^. Results indicated that young SDC4 KO NP and AF have a significant decrease in the ratio of mature-to-immature collagen crosslinks; by 24-months, the ratio of collagen crosslink in NP is higher in KO compared to WT (Fig. 5G). Additionally, peak height ratio of 1549/1338 cm^-1^ (total protein/ collagen) and 1064/1338 cm^-1^ (sulfated GAGs/ collagen) were calculated to determine relative collagen content in tissue. While NP collagen content was relatively similar between genotypes in both age groups, by 24-months of age, SDC4 KO AF had much less collagen content and sulfated GAG (Fig. 5H-I). Notably, sulfated GAG content is significantly higher in young SDC4 KO NP (Fig. 5I). In summary, these findings indicate that SDC4 deletion perturbs the maturation of collagen crosslinking in NP and AF and promotes retention of higher sulfated GAG content in the NP, Altogether, these results provide clear evidence that SDC4 is essential for maintaining the composition and quality of matrix in the intervertebral disc.

**Fig. 5.**
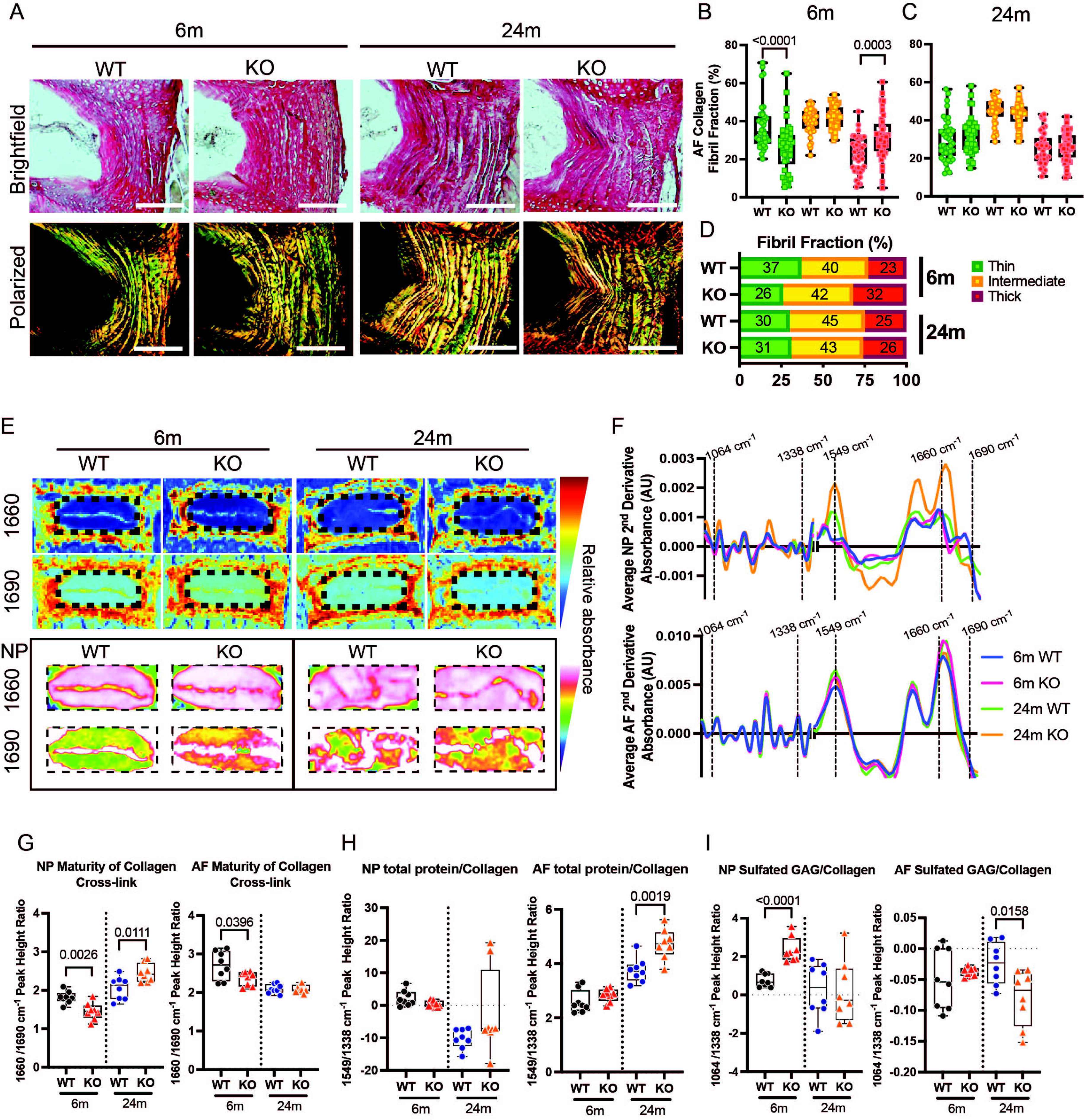
SDC4 deletion alters collagen fiber thickness and matrix composition of the disc. (A) Representative Picrosirius red stained brightfield and polarized images of discs from young adult and old mice. Scale bars, 200µm. (B-D) Quantitative analysis of thin, intermediate, and thick AF collagen fibril fraction. 6-month WT N = 10 animals, n = 40 discs; KO N = 11, n = 42; 24-month WT N = 10, n = 40; KO N = 12, n = 47. (E) Representative chemical map of the disc at 1660cm^-1^ and 1690 cm^-1^ spectra. Zoomed images of NP at these spectra are highlighted to show differences in immature divalent deDHLNL crosslink 1690 cm^-1^ spectrum. The more pink-to-white color shows the higher amount of content detected at peak 1690 cm^-1^. (F) Average second derivative spectra of NP and AF compartments from 6-month and 24-month WT and KO discs were generated, and peak height ratio of (G) collagen maturity (1660/1690 cm^-1^), (H) sulfated GAGs/collagen (1064/1338 cm^-1^), and (I) total protein/collagen (1549/1338 cm^-1^) were calculated. Consequently, the higher the PYR/deDHLNL ratio the more mature collagen cross-links; the smaller the ratio of sulfated GAGs/collagen. N = 4 mice/ genotype, n = 8 discs total per genotype/ timepoint. Quantitative measurements represent the median with the interquartile range. Significance was determined using an unpaired Welch’s t-test or Mann-Whitney test, as appropriate.

### NP matrix composition is altered in SDC4 KO mice

To further ascertain if disc matrix molecules are affected by the absence of SDC4 in the context of disc aging, we performed quantitative immunohistochemistry on young and old SDC4 KO and WT discs. Carbonic anhydrase 3 (CA3), a NP phenotypic marker, was robustly expressed in both genotypes indicating that NP cell health was maintained (Fig. 6A, A’). Previously, we have shown that SDC4 promotes aggrecan turnover, we therefore stained disc sections for chondroitin sulfate (CS), aggrecan (ACAN), and ARGxx, an ADAMTS4/5-cleaved aggrecan neoepitope (Fig. 6B-D). In SDC4 KO, we observed an increase in CS staining, and decreased abundance of ACAN and in ACAN neoepitope ARGxx in AF of young lumbar discs compared to WT (Fig. 6B’-D’). We also determined if SDC4 deletion affects the collagen type I (COL I) and collagen type II (COL II) expression in the AF (Fig. 6E-F) and found that SDC4 KO discs have decreased COL I and increased COL II (Fig. 6E’-F’). These results suggest that deletion SDC4 reduces matrix turnover in young adult lumbar discs evidenced by lower levels of cleaved ACAN neoepitope and an increase in CS expression. In contrast, matrix remodeling may have already occurred by 24-months of age, thus we did not observe further changes in matrix molecules between the genotypes. While there were some differences in COL I and COL II expression, overall result suggests that SDC4 plays a role in maintaining disc tissue homeostasis by maintaining the balance of matrix catabolism and anabolism.

**Fig. 6.**
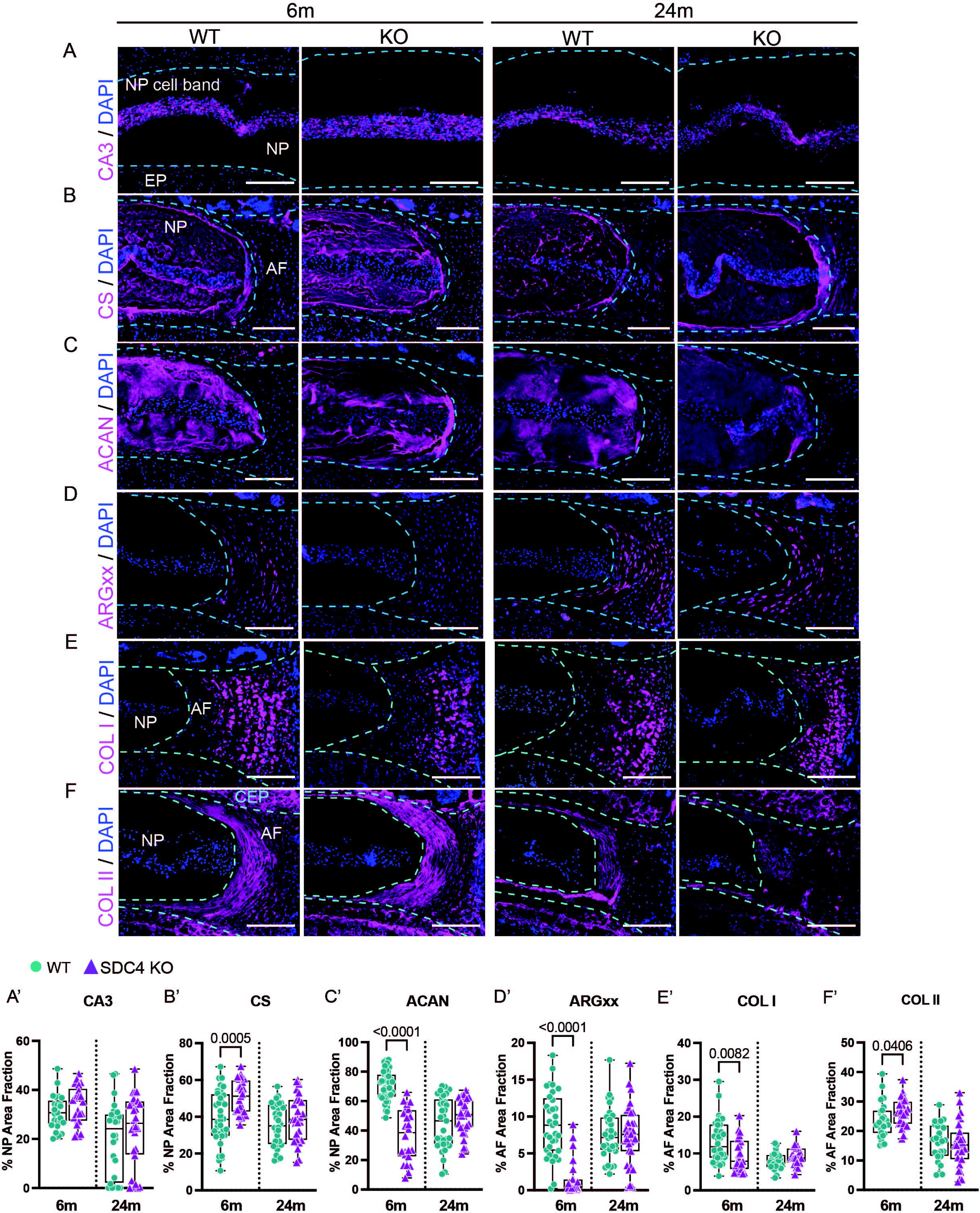
Loss of SDC4 alters NP and AF extracellular matrix composition. Representative images of Immunohistological staining and quantification of (A-A’) NP phenotypical marker CA3, (B-B’) CS, (C-C’) ACAN, (D-D’) ARGxx, (E-E’) COL I, (F-F’) COL II. Scale bars, 200µm. Per marker, 6-month WT N = 7-11 animals, n = 25-43 discs; KO N = 7, n = 24-28; 24-month WT N = 7-8 mice, n = 25-32 discs; KO N = 8, n = 23-32. Dotted lines are used to demarcate NP, AF, and cartilaginous endplate (CEP) compartments. Quantitative measurements represent the median with the interquartile range. Significance was determined using an unpaired Welch’s t-test or Mann-Whitney test, as appropriate.

### Transcriptomic changes reveal prominent dysregulation of heparan sulfate GAG degradation, ER to golgi protein processing, mitochondrial metabolism, and matrix degradation in SDC4 mutant mice

We performed microarray analysis on NP tissue of 6-month-old SDC4 KO and WT discs to identify differentially expressed genes (DEGs) associated with loss of SDC4. Three-dimensional principal component analysis (PCA) showed distinct clusters based on genotypes (Fig. 7A). Analysis of DEGs using False Discovery Rate (FDR) ≤ 5% with a fold change of 2, uncovered 855 downregulated and 290 upregulated genes; these DEGs are represented in a heatmap with hierarchical clustering and volcano plot (Fig. 7B-C).

**Fig. 7.**
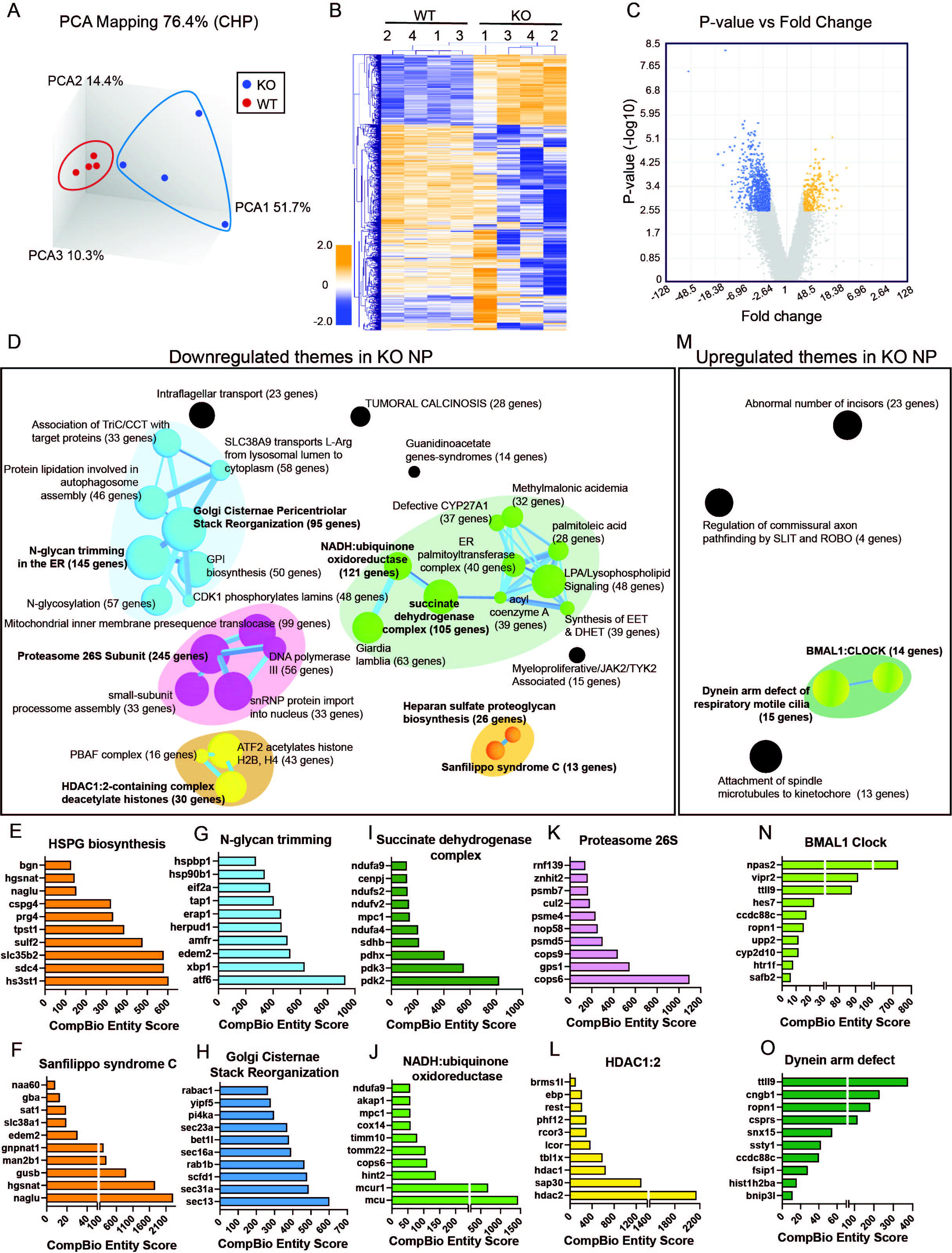
SDC4 KO mice show dysregulation in NP matrix-degrading and secreting processes, metabolic processes, gene regulation and regulation of histone binding. (A) Clustering of transcriptomic profiles by principal component analysis of the NP tissues. N = 4 mice/genotype. (B) Hierarchical clustering of significantly differentially expressed genes (DEGs) (FDR ≤ 0.05, ≥ 2-fold change). (C) Log-log scatterplots of DEGs in the NP. (D) Themes associated with downregulated DEGs are highlighted. The size of a sphere is related to its enrichment score and the thickness of the lines connecting themes signifies the number of genes shared between them. (E-L) CompBio analysis of DEGs and themes whose abundances were significantly downregulated in SDC4 KO NP cells. (M) Themes associated with upregulated DEGs and (N-O) DEGSs and themes whose abundances were significantly upregulated.

To gain insights into possible biological features associated with SDC4 loss-of-function, CompBio tool (PercayAI Inc., St. Louis, MO) was used. CompBio tool assembled contextually relevant information by performing a literature analysis on SDC4 KO DEGs to identify relevant concepts and pathways and present them as themes (colored spheres) and clusters of related themes (colored spheres and sticks). In downregulated themes (Fig. 7D), five superclusters were identified: 1) heparan sulfate proteoglycan biosynthesis, 2) N-glycan trimming in endoplasmic reticulum (ER) and golgi cisternae stack reorganization, 3) mitochondria machinery, 4) proteasome particle regulatory, and 5) transcription regulation. Of these superclusters, the top themes with the highest CompBio normalized enrichment score (NEScore) were highlighted (Fig. 7E-L, N-O). SDC4 deletion significantly affected genes relevant to the heparan sulfate biosynthetic enzymes, such as *hs3st1*, *slc35b2*, *sulf2*, *cspg4*, *prg4*, *bgn* (Fig. 7E). As expected, *sdc4* which showed the highest fold change of 54-fold was also enriched under this theme. Notably, each gene was previously reported to be involved in sulfation of GAGs and extracellular matrix homeostasis(21–26). Additionally, DEGs, such as *naglu*, *hgsnat*, *gusb*, *man2b1*, were contextually grouped and identified as a theme relating to Sanfilippo syndrome C, a lysosomal disease relating to catabolic enzymes that degrade heparan sulfate(27–29) (Fig. 7F). Themes such as Defective CYP27A1 and Synthesis of epoxy (EET) and dihydroxyeicosatrienoic acids (DHET) themes also suggest decrease of autophagy and matrix degradation (Supp. Fig. 1A-S). In the second supercluster, downregulated genes relating to ER-golgi protein trafficking, and golgi cisternae stack reorganization include *atf6*, *xbp1*, *edem2*, *sec13*, *sec31a*, *scfd1*, *rab1b* (Fig. 7G-H). This suggests impaired ER-mitochondria calcium homeostasis, autophagy, and adaption to variations in ER cargo load or to the accumulation of glycoprotein in NP cells(30–32). Additionally, in the third supercluster, themes related to mitochondria machinery and processes involving the succinate dehydrogenase complex and NADH ubiquinone oxidoreductase show downregulation of *mcu*, *mcur1*, *pdk2*, *pdk3*, *pdhx* (Fig. 7I-J) suggesting altered cellular bioenergetics (33). In the proteasome particle regulatory supercluster, genes include *cops6*, *gps1*, *cops9*, and *psmd5* (Fig. 7K). Lastly, genes grouped in the transcription regulation supercluster include *hdac1*, *hdac2*, *sap30*, *tbl1x* (Fig. 7L). All superclusters in NP downregulation themes suggest perturbation in matrix breakdown, protein trafficking, mitochondria-autophagosomes and transcription regulation.

Of the 290 upregulated genes, only 54 genes contributed to 4 enriched themes meeting a statistical significance cutoff. A mini cluster of themes was identified with BMAL1 Clock regulation (Fig. 7M). Here, there are 2 themes, one of which is BMAL1; it is mainly driven by *npas2*, a substitute for BMAL1 that maintains circadian behaviors(34) (Fig. 7N). The other theme relates to the dynein arm defect of cilia, driven by *ttll9* (Fig. 7O). Taken together, these themes suggest SDC4 deletion affects NP cell cilia and some components of the NP circadian clock. The overall contribution of SDC4 to the homeostasis of the spinal motion segment comprising vertebrae and intervertebral disc is summarized in Figure 8.

**Fig. 8.**
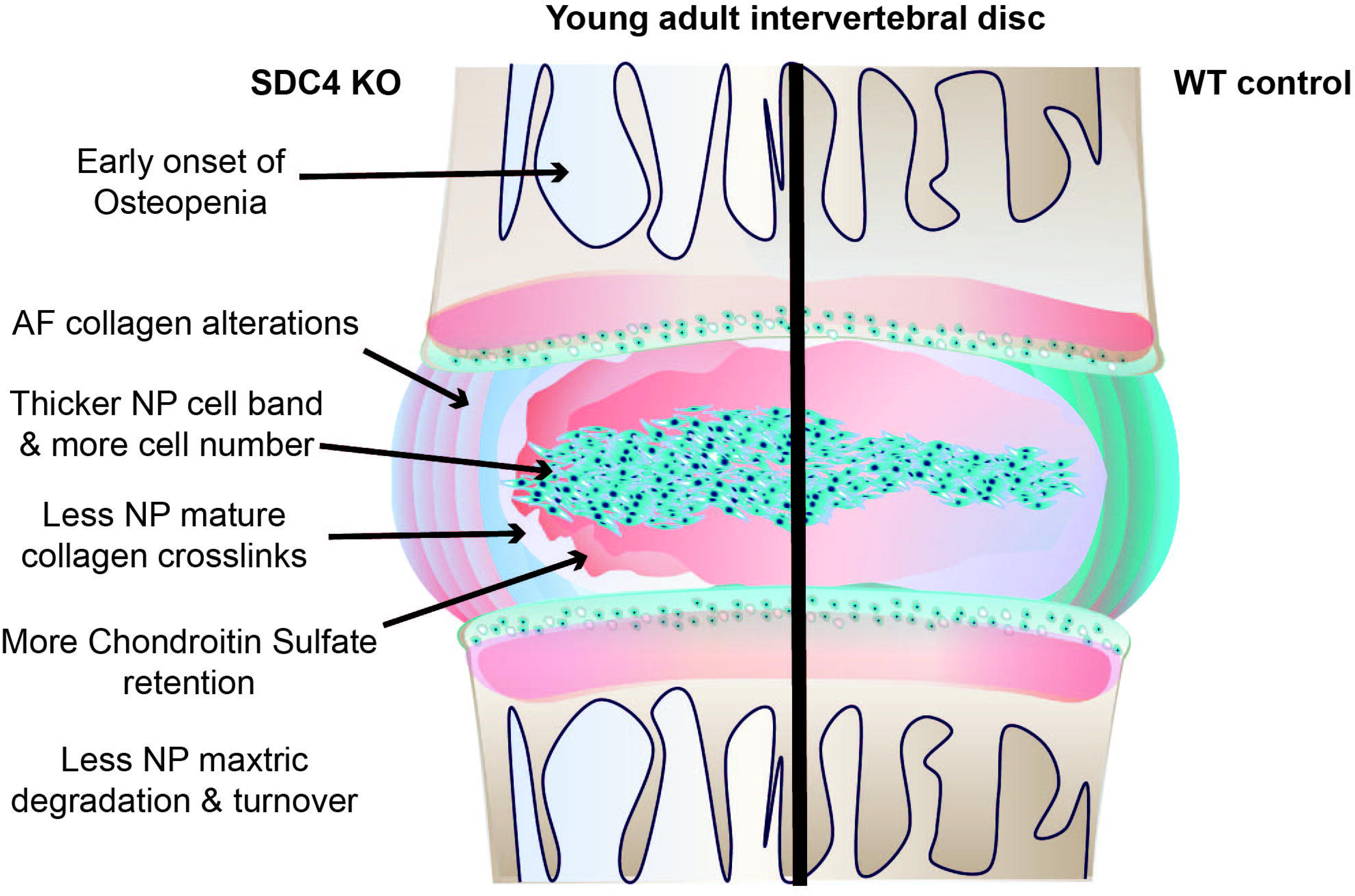
An illustration of alterations in matrix homeostasis and vertebral bone geometry of the intervertebral disc and spine induced by loss of SDC4 function.

## Discussion

Our findings show for the first time that SDC4 deletion instigates early onset of vertebral osteopenia in mice. We show severe alterations in trabecular bone geometry and cortical bone thinning in adult mice which progressively worsens with age. This alteration also includes reduced bone mineral density, tissue mineral density, and microarchitectural changes in the trabecular tissue. Mechanistically, we determined that the changes in vertebral bone structure were not due to intrinsic osteoblastic defects in mineral deposition but rather elevated osteoclastic activity. This increase in osteoclastic activity may be due to the absence of heparan sulfate from SDC4 deficiency. In support of this idea, Kim et al. detailed the inhibitory effect of heparan sulfate on osteoclast differentiation and demonstrated that in mice periosteal regions of calvaria injected with syndecan ectodomains the formation of TRAP-positive mature osteoclasts on the calvarial bone surface was reduced(35). Altogether, these structural deficits contribute to lower stiffness and resistance to deformation in vertebral bones of SDC4 KO mice.

Histological assessments of middle and old-aged discs were comparable between genotypes. However, young SDC4 KO discs possessed a relatively larger sized cell band and higher cell count compared to their age-matched controls. Given the alterations in bone mineral density and biomechanics, it is plausible that KO discs at this age experience lower loads and thus exhibit slightly larger NP cell vacuoles, cell bands, and cell numbers. For example, Bian et al. demonstrated a reduction of NP vacuoles under mechanical stress(36). Because NP cells experience osmotic and mechanical loading, we considered whether SDC4 deletion results in altered mechanosensing in these cells(37). However, our observations suggest that SDC4 is not essential in mediating NP cell mechanosensing.

The maintenance of a disc hydrophilic milieu during spinal loading depends on ECM quality and the homeostatic balance between matrix anabolism and catabolism. We previously detailed the contribution of SDC4-ADAMTS5 axis in aggrecan degradation(13, 16). Since SDC4 KO revealed subtle phenotypic differences, imaging-FTIR was used to better capture the chemical signatures of these changes. In addition to the reduced COL I staining in the AF, spectroscopy revealed that discs of young adult SDC4 KO mice have a relatively lower ratio of mature collagen crosslinking in NP and AF compartments compared to WT suggesting SDC4’s involvement in mediating collagen type I secretion and crosslinking. Other studies showing SDC4 affecting collagen type I production and crosslinking in renal fibrosis and collagen crosslinking in the heart support our observations(38, 39). Interestingly, old KO NP has a higher relative ratio of mature collagen crosslinking compared to WT. This accumulation could be explained by the decreased matrix turnover in young SDC4 KO, evidenced by ARGxx staining. Consistent with this, decreased thin and increased thick collagen fibrils in young SDC4 KO may suggest a decline in KO collagen turnover. Moreover, in young lumbar discs, SDC4 deficiency resulted in increases in chondroitin sulfated GAGs and collagen II compared to WT. This is in accordance with previous reports that SDC4 deletion reduces collagen I and fragmented aggrecan neoepitope, ARGxx, implying diminished proteolytic cleavage of the matrix(13, 16, 38).

Further complimentary insights concerning SDC4’s broader role in the disc came from transcriptomic data analysis. SDC4 deletion affected the downregulation of genes pertaining to 5 major themes. Our data shows themes critical in heparan sulfate synthesis and Sanfilippo syndrome C, which contain genes involved in GAG sulfation and lysosomal degradation of large sugar molecules such as heparan sulfate. Their downregulation likely resulted in the accumulation of sugars like heparan sulfate causing deleterious effects on tissue homeostasis in rare diseases such as Mucopolysaccharidosis(40). Similarly, themes relating to DNA transcription and ER-protein folding and trafficking were identified. This includes the Golgi Cisternae Stack reorganization theme containing a high enrichment of *sec13*, *sec23a*, and *sec31a*, components of coat protein complex type II (COPII) involved in procollagen trafficking. *sec13* downregulation can cause defects in collagen secretion; this corresponds with the lower collagen I expression observed in SDC4 KO AF(41). Interestingly, downregulation of *sec23a* and *sec31a* is associated with skeletal defects and delayed growth(42). Furthermore, low levels of *pdhx, pdk2, pdk3, mpc1, mcu,* and *mcur1* suggest NP cells maintain bioenergetics in the hypoxic tissue through shunting of glucose metabolites from the mitochondria to glycolysis to sustain ATP production and attenuate hypoxic reactive oxygen species generation(43–46). Low levels of *mcu* and *mcur1* may also help maintain low levels of calcium in the mitochondrial matrix protecting against apoptosis(47). In the Defective CYP27A1 and Synthesis of EET and DHET themes, downregulation of *naaa* and *nampt* suggests a decrease of autophagy and matrix degradation(48) bolstering the observed preservation of NP cell health and decrease in matrix catabolism and matrix turnover. Additional clusters include 26S proteasome and transcription regulation themes containing genes that regulate cell cycle. In SDC4 KO NP upregulated themes, two were identified based on their statistical cutoff and normalized enrichment score. Themes relating to Cilia and BMAL1 Clock include upregulation of *ttl9* and *npas2*, respectively. *Bmal ^-/-^* mice have exhibited premature aging phenotypes and disc degeneration suggesting its importance in tissue health maintenance(49, 50), and it has been suggested that *npas2* can serve as a functional substitute(51). Consequently, upregulation of *npas2* suggests loss of SDC4 may preserve NP matrix.

In summary, our study provides insights into the role of SDC4 in vertebral bone health and disc matrix homeostasis (Fig. 8). SDC4 loss promotes early onset of osteopenia and alterations in vertebral bone biomechanics which could alter spinal loading and load dissipation. SDC4 loss also resulted in reduction in matrix degrading activity, a relatively lower ratio of mature collagen crosslinking, and an increase in chondroitin sulfate GAGs which point to its role in preserving NP health in young mice. However, CompBio analysis highlighted that these observations are likely to be secondary effects from downregulated lysosomal degradation and collagen secretion, resulting in GAGs accumulation and collagen I decrease. In summary, our work shows loss of SDC4 may be beneficial in certain contexts, however, SDC4 is still necessary for spinal health, particularly in the maintenance of bone and matrix secretion in the intervertebral disc.

## MATERIALS AND METHODS

### Mice

All animal care procedures, housing, breeding, and the collection of animal tissues were performed in accordance with a protocol approved by the Institutional Animal Care and Use Committee (IACUC) of Thomas Jefferson University. The global SDC4 knockout mice on C57BL/6 background have been described earlier and obtained from Dr. Michael Simons(11, 52). These mice harbor deletions of exon 2 to part of exon 5 that encodes the N-terminal coding region. Both male and female mice were used in these studies.

### Micro-Computed Tomography (µCT) Analysis

SDC4 KO and WT lumbar spines were dissected and fixed in 4% PFA in PBS at 4°C for 48 hours prior to µCT scans (Bruker, Skyscan 1275). For assessment of bone morphometry, the lumbar spines of SDC4 KO and WT mice were rinsed and hydrated in 1x PBS, scanned at a resolution of 15 µm^3^ voxel (50 kV, 200 µA, 85 ms exposure time, rotation step of 0.2°, using 1 mm aluminum filter). The reconstruction was performed using Skyscan NRecon package. Distance measurements were made along the dorsal, midline, and ventral regions in the sagittal plane of disc height and vertebral length and then averaged, which was then used to calculate the disc height index (DHI), as previously described(53, 54). Trabecular parameters were measured using Skyscan CT analysis (CTAn) software by contouring the region of interest (ROI) in the 3D reconstructed trabecular tissue. Resulting datasets were assessed for bone volume fraction (BV/TV), trabecular number (Tb. N.), trabecular thickness (Tb. Th.), and trabecular separation (Tb. Sp.). The cortical bone was analyzed in two dimensions and assessed for bone volume (BV), cross-sectional thickness (Cs. Th.), mean cross-sectional bone area (B. Ar), and mean cross-sectional tissue area (T. Ar). Mineral density was calculated using a standard curve created with a mineral density calibration phantom pair (0.25 g/cm^3^ calcium hydroxyapatite (CaHA) and 0.75 g/cm^3^ CaHA)).

### Biomechanical Testing

Lumbar vertebrae (L1-L6) from 6-month-old WT and SDC4 KO mice were isolated, removed of cartilaginous endplates and articular processes, and µCT scanned. CT images were used to determine the vertebral diameter and average height of each vertebra. Samples underwent not more than two freeze-thaw cycles prior to testing. Each vertebra was individually potted into a 2-mm plastic ring mold using an acrylic resin (Ortho-Jet, Patterson Dental), and mechanical loading was applied using a material testing system with a 50 lb load cell (TA Systems Electrofoce 3200 Series II). A 0.4 N compressive preload was applied, followed by a monotonic displacement ramp at 0.1 mm/s until failure. Force-displacement data were digitally captured at 25 Hz and converted to stress-strain using a custom GNU Octave script with µCT-based geometric measurements, as previously described(55, 56).

### TRAP and TNAP Staining

Frozen sections of fixed WT and SDC4 KO lumbar spines (L1-6) were cut at 10 µm onto cryofilm to maintain the morphology of the mineralized sections. The taped sections were glued to microscope slides using chitosan adhesive, dried overnight at 4°C, and rinsed with PBS prior to staining. For TNAP staining, slides were incubated for 10 min in 100mM Tris-HCl buffer (AP Buffer) (pH 8-8.5) mixed with Vector Blue (Vector laboratories, SK-5300) at room temperature following the manufacturer’s instructions. Subsequently, samples were rinsed in PBS for 5 min 3 times, mounted with DAPI and visualized under Cy5 channel. For TRAP staining, ELF97 (ThermoFisher Scientific, E6588) was diluted 1:75 in TRAP buffer (9.2 g of sodium acetate anhydrous and 11.4 g of sodium tartrate dibasic dihydrate dissolved in 1L water; pH 4.2). TRAP buffer was applied to samples for 15 min at room temperature. Subsequently, ELF97-diluted TRAP buffer was applied to the slides for 5 min in room temperature. The slides were rinsed in PBS for 5 min 3 times, mounted with DAPI, and visualized under the GFP channel.

### Osteoclast and Osteoblast Cultures

Bone marrow was pooled from long bones of 2-4-month-old mice (3 animals/genotype per experiment) by removing the ends of the bones to expose the bone marrow cavity and flushed using 21G syringe. Pooled bone marrow stromal cells (BMSCs) were washed with 3 times with PBS and cultured in αMEM (Sigma, M2279) supplemented with 20% FBS, Penicillin/Streptomycin, 2 mM Glutamine. After 2 days post-isolation, non-adherent hematopoietic progenitors were pooled and plated at 7x10^4^ cell/well (96-well) with complete media supplemented with 30 ng/mL M-CSF (R&D Systems, 416-ML). At 90% confluence, osteoclast differentiation was initiated using αMEM complete media supplemented with 30 ng/mL M-CSF and 100 ng/mL RANKL (R&D Systems, 462-TEC/CF)(57). 50% osteoclast differentiation medium was changed every other day, for 6-9 days. Differentiated cells were fixed with 4% PFA and stained with TRAP staining buffer as described(57). Cells were imaged and TRAP-positive osteoclasts with 2 or more nuclei per cell body were counted.

For osteoblast culture, expanded adherent BMSCs were plated in 96 well plates at confluence before adding osteogenic medium: DMEM (Corning) complete media supplemented with 10 mM β-glycerophosphate (Sigma, G9422) and 50 µg/mL L-ascorbicacid-2-phosphate (Sigma, 49752)(57). 50% osteogenic medium was changed every other day, for 14 days. Cells were fixed and stained with TNAP (Vector laboratories, SK-5300) following manufacturer instruction and alizarin red as described to detect the presence of calcium deposits, respectively.

### Histological Analysis

Lumbar spines were dissected and immediately fixed in 4% PFA in PBS at 4°C for 48 hours, decalcified in 20% EDTA at 4°C for 2-3 weeks, and then embedded in paraffin. 7 µm mid-coronal sections were cut from L3-S1 levels and used for histological staining (N = 7-11 animals per genotype/time point, n = 23-43 discs/genotype). Sectioned tissues were deparaffinized, followed by graded ethanol rehydration preceded all staining protocols. Following Safranin-O/Fast Green/Hematoxylin staining, images were acquired on a light microscope (Axio Imager 2; Carl Zeiss Microscopy) using 5x/0.15 N-Achroplan (Carl Zeiss) objective and Zen2™ software (Carl Zeiss). The health of disc compartments was assessed by at least four blinded graders using Modified Thompson Grading(56, 58–61). Picrosirius red staining (Polysciences, 24901) was performed to assess collagen fibril thickness, and images were acquired using 4x/0.25 Pol /WD 7.0 (Nikon) objective on a polarizing light microscope (Eclipse LV100 POL; Nikon). NIS Elements Viewer software (Nikon) was used to set color threshold for the level of green (thin), yellow (intermediate), and red (thick) fibers(56, 60). Color threshold levels remained constant for all samples.

### Imaging Fourier-Transform Infrared Spectroscopy

7 µm deparaffinized sections of decalcified lumbar disc tissues (L5/S1) were used to acquire infrared (IR) spectral imaging data using methods previously described(56, 60). Spatial resolution images were acquired on the Spectrum Spotlight 400 FT-IR Imaging system (Perkin Elmer, Waltham, MA, USA) in the mid-IR (MIR) region from 4000 cm^−1^ to 740 cm^−1^ wavenumbers spectral range, at 25 µm pixel resolution, 8 scans per pixel, and a spectral resolution of 8 cm^-1^. Spectra were collected across the MIR region of three consecutive sections per tissue sample. MIR spectral imaging data were analyzed using ISys Chemical Imaging Analysis software (v. 5.0.0.14). Raw spectra went through an atmospheric correction and converted to second derivative spectra to resolve underlying peaks and corrected for baseline offsets. Savitsky-Golay smoothing (filter order of 3 and filter length of 9) was applied, and spectra inverted by multiplication by -1 to convert to positive peak heights as appropriate(56, 60). Mean second-derivative absorbances in the amide I (1660 cm^-1^), amide II (1549 cm^-1^), collagen side chain vibration (1338 cm^-1^), and sulfated proteoglycan sugar ring (1064 cm^-1^) peaks were quantified and compared in the NP and AF compartments. The ratio of collagen maturity (1660/1690 cm^-1^), sulfated GAGs/collagen (1064/1338 cm^-1^), and total protein/collagen (1549/1338 cm^-1^) were calculated(20) using peak heights derived from mean second-derivative absorbance.

### Immunohistochemistry and digital image analysis

Mid-coronal disc sections (7 μm) were deparaffinized in histoclear and rehydrated in ethanol solutions (100–95%), water, and PBS. Citrate-buffer at pH 6 (Vector Laboratories) antigen retrieval method was performed on samples required for collagen 1 (1:100; Abcam, ab34710), collagen II (1:400; Fitzgerald, 70R-CR008), CA3 (1:150; Santa Cruz Biotechnology, sc-50715), CS (1:300; Abcam, ab11570); 1:500 proteinase-K enzymatic retrieval method for ARGxx (1:200; Abcam, ab3773); 1:200 chondroitinase ABC (20 U/mL) enzymatic retrieval for ACAN (1:50; Millipore Sigma, AB1031). MOM kit (Vector Laboratories; BMK-2202) was used per manufacturer instruction. After antigen retrieval, samples were blocked with 5-10% normal goat or donkey serum (Jackson ImmunoResearch) in PBS-Triton (0.4% Triton X-100 in 1x PBS) for 1 h at room temperature. Primary antibodies were applied and incubated overnight at 4°C.

Samples were then generously washed three times with PBS and incubated with Alexa Fluor-594-conjugated secondary antibody (1:700, Jackson ImmunoResearch) for 1 h at room temperature, shielded from light. Samples were washed three times with PBS and mounted with ProLong Gold Antifade Mountant with DAPI (Thermo Fisher Scientific, P36934). Images were acquired on Axio Imager 2 (Carl Zeiss Microscopy) using 5×/0.15 N-Achroplan (Carl Zeiss Microscopy) or 10×/0.3 EC Plan-Neofluar (Carl Zeiss Microscopy) objective, X-Cite 120Q Excitation Light Source (Excelitas Technologies), AxioCam MRm R3 camera (Carl Zeiss Microscopy), and Zen 2^TM^ software (Carl Zeiss Microscopy). Per staining experiment, images were taken at a set exposure time for all samples. All quantifications were done on ImageJ 1.52i (NIH). Images were thresholded to create binary images, and NP and AF compartments were manually contoured using the Freehand Tool. These ROI were analyzed using either the Analyze Particles (cell number quantification) function or the Area Fraction measurement. Cell area and cell perimeter were measured using the Freehand Lines Tool. Cell numbers per disc were calculated from averaged DAPI-positive cells of three independent tissue sections of a disc.

### Mouse NP cell isolation and staining

NP tissues from three mice per genotype were pooled and scooped into low glucose DMEM (Corning) supplemented with 10% FBS and Penicillin/Streptomycin. Tissues were gently washed 3 times with PBS with Pen/Strep and centrifuged at 800 rpm for 3 minutes. Tissues were digested with 2 mg/mL collagenase P (Sigma) for about 10 min, neutralized, and washed with complete media 3 times at 800 rpm for 3 minutes. Cells were resuspended and plated onto 10 µg/mL fibronectin- or rat tail collagen type I-coated glass bottom dishes (Sigma). Cells were fixed the next day with 4% paraformaldehyde, permeabilized with 0.25% triton-x and stained with p-Paxillin (CST, 69363) and YAP (Santa Cruz Biotechnology, sc-101199). Images were captured on Zeiss LSM 800 Axio Inverted confocal microscope (Plan-Apochromat 63x/1.40 oil).

### Tissue RNA isolation

NP tissues from lumbar spines (L1/2-L6/S1) were micro-dissected from 6-month-old WT and SDC4 KO mice (n=4 mice/genotype, WT = 3 F, 1 M; KO = 2 M, 2 F) using a dissecting microscope (Zeiss, Stemi 503). Each sample consisting of 6 lumbar NP tissues was pooled and stored in RNAlater® Reagent (Invitrogen) at 80°C until isolation. Tissues were homogenized with a Pellet Pestle Motor (Sigma Aldrich, Z359971), and RNA was extracted using RNeasy Micro Kit (Qiagen, 74004) following manufacturer instruction.

### Microarray analysis

Total RNA integrity was assessed on Agilent 4150 TapeStation. Samples with RIN > 5 were used for microarray. GeneChip™ Whole-Transcriptome (WT) Pico Kit was used per manufacturer instruction. Briefly, the kit was used to amplify coding RNA from 2 ng total RNA and subsequently generate single-strand cDNA, which were then fragmented and labeled. The fragmented and biotin-labeled ss-cDNA were hybridized to mouse Clariom™ S Assay cartridges overnight. The cartridges were wash and stain with GeneChip™ hybridization wash and stain kit, and then scanned on the Affymetrix GeneChip™ Scanner 3000 7G System. Raw data were processed with Transcriptome Analysis Console (TAC) v 4.0.2 (Applied Biosystem). Gene summarization using the Signal Space Transformation-Robust Multi-Array Average (SST-RMA) algorithm was first applied to summarize, background subtraction, and normalize the CEL files to generate CHP files. These files were then used to carry out differential gene expression analysis using ANOVA method for statistical testing with ebayes (Empirical Bayes Statistics for Differential Expression) correction for small sample size and filtered for genes that are ≥ 50% detected above background (DABG). Significant results were considered for transcript that showed a fold change of ±2 with False Discovery Rate (FDR) ≤ 0.05. The array data are deposited in a publicly accessible GEO database (GSE243573).

### Transcriptomic data analyses using CompBio tool

Significantly up- and downregulated DEGs from NP tissues of SDC4 mice (A 2-fold difference, FDR < 0.05) were analyzed using the GTAC-CompBio Analysis Tool (PercayAI Inc., St. Louis, MO). CompBio uses automated Biological Knowledge Generation Engine (BKGE) to extract all abstracts from PubMed that reference the input DEGs to identify relevant processes and pathways(46, 61, 62). Conditional probability analysis is used to compute the statistical enrichment score of biological concepts (processes/pathways) over those that occur by random sampling. The scores are then normalized for significance empirically over large, randomized query group. The reported normalized enrichment scores (NEScore) represent the magnitude to which the concepts/themes are enriched above random, and an empirically derived p-value identifies the likelihood of achieving that NES by chance. The overall p-value of ≥ 0.1 with NEScore of ≥ 1.2 are used, resultant up- and downregulated thematic matrices are presented.

### Statistical analysis

Statistical analysis was performed using Prism 10 (GraphPad, La Jolla, CA, USA) with data presented as whisker box plots showing all data points with median and interquartile range and maximum and minimum values. Differences between distributions were checked for normality using Shapiro–Wilk tests and further analyzed using an unpaired Welch’s t-test for normally distributed data and the Mann–Whitney U test for non-normally distributed data. Analyses of Modified Thompson Grading data distributions and fiber thickness distributions were performed using a chi-square test at a 0.05 level of significance.

## DATA AVAILABILITY

RNA microarray data associated with this study are deposited in the GEO database (GSE243573). All datasets generated and analyzed during this study are included in this published article.

## ACKNOWLEDGMENTS

We would like to thank Alexandra Ciuciu and Dr. Ryan Tomlinson for their helpful suggestions on biomechanics study and Dr. Andrzej Steplewski, Thomas Jefferson University for the FTIR spectroscopy. We also thank Dr. Ruteja Barve for her excellent technical assistance on CompBio.

## CONFLICT OF INTERSTS

Authors of this manuscript do not have conflicts of interest to disclose.

## AUTHOR CONTRIBUTIONS

K.S and M.V.R. conceived and designed the experiments. K.S. performed the experiments, collected, and analyzed the data. K.S. and M.V.R. interpreted the data. K.S. and M.V.R. drafted the manuscript.

## ETHICS STATEMENT

All animal experiments were performed under IACUC protocols approved by Thomas Jefferson University.

