## Supplementary figures and images for "SDC4 deletion perturbs intervertebral disc matrix homeostasis and promotes early osteopenia in the aging spine"

### Supplemental Fig 1

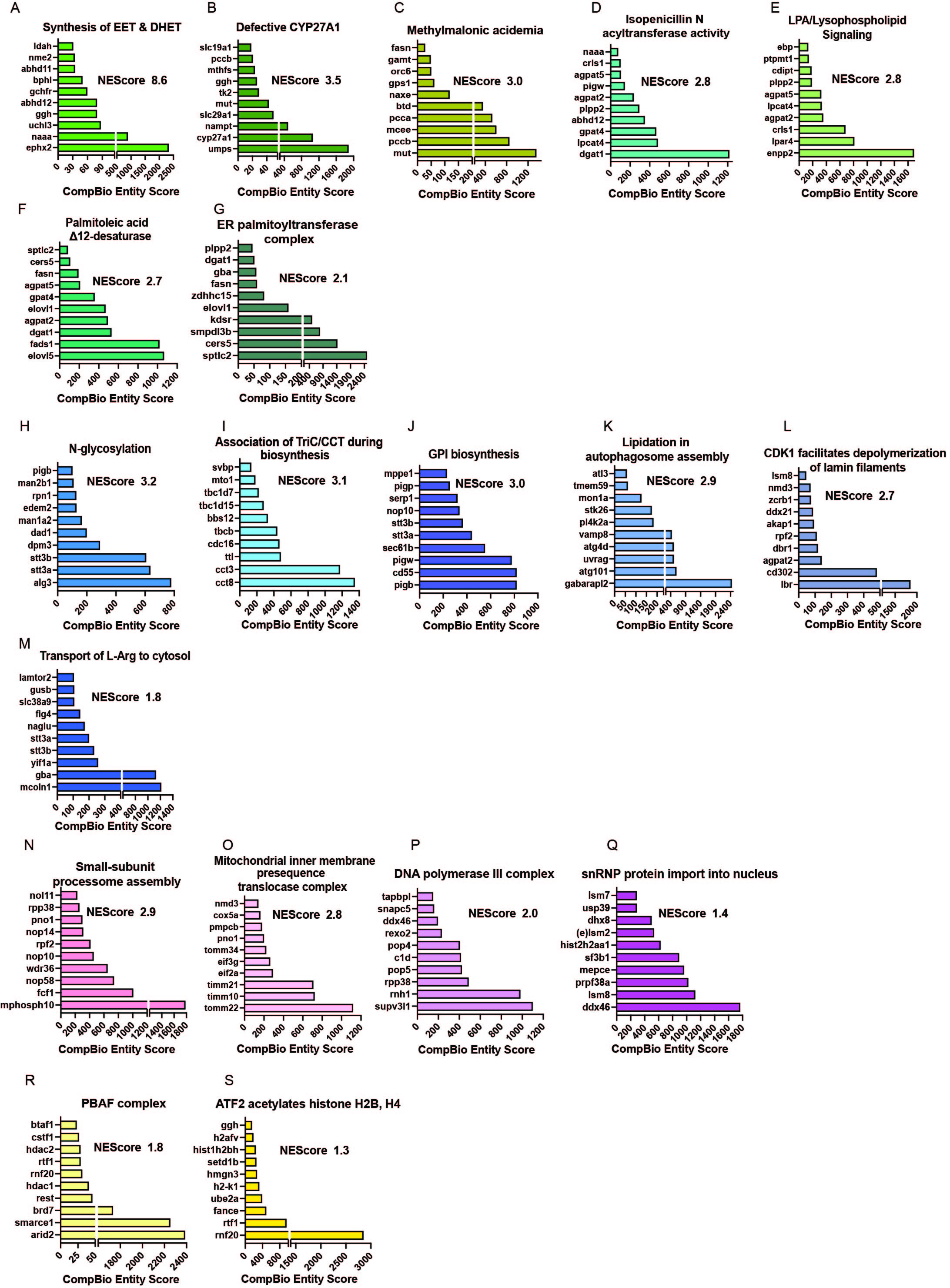
